# Connecting the dots - Recognition of artificial and natural shapes relies on representing points of high information

**DOI:** 10.1101/2025.11.17.688832

**Authors:** Gunnar Schmidtmann, Nicholas Baker, Kevin J. Lande, Filipp Schmidt

## Abstract

Physiological and psychophysical evidence suggests that the visual system represents object outlines using prominent curvature features, particularly regions of extreme curvature (convex maxima and concave minima). These curvature extrema often coincide with points of high informational content (“surprisal”), but this relationship is only correlational. It remains unclear whether the visual system explicitly encodes curvature extrema or instead prioritizes the most informative contour locations. To address this, we conducted two shape-matching experiments comparing the roles of curvature extrema and surprisal in shape representation. Observers performed match-to-sample tasks in which smooth reference shapes were matched to simplified polygonal versions created by connecting subsets of contour points corresponding to (i) curvature maxima, (ii) curvature maxima and minima, or (iii) points of highest surprisal. Stimuli included artificial shapes composed of compound radial frequency patterns and natural shapes (animal outlines), the latter allowing us to dissociate curvature and information by restricting sampled points. Performance was higher for natural than artificial shapes (95% vs. ∼86%). Shapes defined by a small number of high-surprisal points matched performance in baseline and curvature maxima and minima conditions, but exceeded performance for curvature maxima alone (∼90% vs. ∼65%). In a second experiment, we contrasted surprisal with curvature maxima and minima conditions while varying the number of sampled points (4–32). Performance increased with point number, approaching ∼90%. Critically, under strong simplification (4–6 points), surprisal-based shapes yielded higher accuracy than curvature-based shapes. These findings suggest that shape representation emphasizes features with high informational content rather than curvature extrema per se.

## Introduction

Object and shape recognition is a fundamental aspect of human vision. Despite huge variations in the structure and form of objects and their retinal projections, humans recognize shapes quickly and accurately (see recent review by Elder, 2018). While objects in the visual world are usually three-dimensional (3D), our retinas process visual information based on a two-dimensional (2D) projection. Furthermore, each 3D object can create an infinite number of projections on the retina, depending on object position (viewing distance), scale, and countless spatial transformations (e.g., pose, rotation, slant, etc.). Human object recognition is remarkably robust to these variations (DiCarlo et al., 2012; Elder, 2018). This is also exemplified by the fact that humans recognize objects from images on planar surfaces, like pictures, photographs, or screens (e.g., television). Note that object recognition refers to the ability to identify objects across changes in appearance resulting from differences in viewpoint, scale, etc., whereas shape recognition refers to the extraction and representation of structural form information that supports this invariance. Our focus on planar shapes is motivated by the need to understand how shape-based representations enable successful object recognition even without depth cues or surface information.

While recent developments in computer vision and machine learning have overcome some of the challenges associated with human perceptual invariance in object recognition, human-like performance, which is largely independent of spatial transformation, continues to be a challenge for computer vision (e.g., Kar & DiCarlo, 2024; Pinto et al., 2008; Rajalingham et al., 2018).

Therefore, an important question in vision science is how the human visual system represents the vast variety of planar shapes to achieve perceptual invariance. Throughout the long history of research on shape perception, different theories have been offered as to how shapes are represented—whether in terms of curvature, medial axis skeletons (Blum, 1967, 1973; Feldman & Singh, 2006), shock graphs (Rezanejad & Siddiqi, 2013), feature vectors, or otherwise (see Elder, 2018 and Todd & Petrov, 2022 for reviews).

This study aims to investigate the hypothesis that shape recognition is heavily influenced by the specific curvature features along the contour of a closed shape—namely, points of positive curvature maxima (convexities, or corners and bumps) and points of negative curvature minima (concavities, or creases and dents).^1^ Our findings confirm that both convexities and concavities have a significant role in object recognition. In addition, we investigate the psychological relationship between these curvature features and information content. Curvature extrema in general, and concave minima especially, have been shown to correlate with points of high information along the contour (Resnikoff, 1985; Feldman and Singh, 2005). In other words, areas of extreme curvature are areas in which contour changes are least predictable. One hypothesis is that the visual system encodes curvature extrema because they are effective proxies for informative points along a contour. However, another possibility is that the visual system directly extracts points of high information content, which just happen regularly to coincide with curvature extrema. Our findings provide initial support for the latter hypothesis.

### Importance of regions of high curvature

While the existing literature has pointed to the critical importance of curvature in shape processing, there is little consensus on which specific features of curvature are significant, with results often depending on the experimental task and the stimulus design. Attneave (1954) demonstrated the significant role of regions of *high curvature* in shape recognition by isolating the points of maximum and minimum local curvature (convexities and concavities) in an image of a sleeping cat and connecting them with straight lines, thereby omitting intermediate shape information. Despite the absence of this intermediate contour information, the sleeping cat was still recognizable (for a generalization see De Winter & Wagemans, 2008a). In another experiment (never fully published), Attneave showed that when observers are asked to indicate the most salient points along a smooth closed shape, they consistently selected convexities and concavities. The latter results were more recently confirmed by Norman et al. (2001). They demonstrated that when observers are presented with silhouettes of smooth curved shapes (sweet potatoes) and asked to locate on the outline ten points that they considered the most informative, observers consistently chose regions of high curvature of both signs (convexities and concavities). The same has been shown for line drawings of everyday objects (De Winter & Wagemans, 2008b). Further evidence for the importance of both convex and concave curvature comes from studies investigating change detection (Bertamini, 2008), visual search (Bertamini & Lawson, 2008), and shape adaptation (Bell et al., 2010).

### Comparing the roles of convexities and concavities

Many studies have found evidence that among points of high curvature, convexities (i.e. bumps) are more crucial for shape recognition than concavities (i.e., dents). For instance, Biederman (1987) demonstrated that removing regions of maximum convex curvature from images of familiar shapes impaired recognition more significantly than removing concave or intermediate regions. Schmidtmann et al. (2015) conducted experiments where observers had to match a segmented shape (based on compound radial frequency (RF) patterns, see Methods below) showing either regions of maximum convexities, concavities or intermediate regions of varying segment lengths (ranging from single points to longer segments) to one of two subsequently presented whole-contour shapes. Their results show that for very short segment lengths, performance was significantly better for convexities compared to concavities or intermediate regions and was independent of segment length. For concavities and intermediate points, on the other hand, performance was at chance-level for very short segments, and improved with increasing segment length, but only reached the same performance as convexities for long segments. Schmidtmann et al. (2015) concluded that for this shape-matching experiment, closed curvilinear shapes are encoded using the positions of convexities, rather than concavities or intermediate regions (see Discussion for further details).

A similar advantage for convex points has been demonstrated in shape discrimination experiments. Loffler et al. (2003) measured shape discrimination between RF patterns and circles and showed that sensitivity was significantly impaired when points of maximum convex curvature were occluded. Likewise, in a lateral masking paradigm, Poirier and Wilson (2007) found that convex masks had a greater influence on the discrimination of RF patterns than minimally curved (i.e., flat-ish) sides. Other studies show that convexity has a privileged role in the extraction of other features, for example in positional discrimination (Bertamini, 2001; Bertamini & Farrant, 2006; Gibson, 1994) or the detection of symmetry (Hulleman & Olivers, 2007), though the latter might be caused by the tendency to attend to convexities features (Bertamini et al., 2013).

Comparatively fewer studies have reported an advantage for concave features. Barenholtz et al. (2003) measured detection of changes in shapes and reported an advantage for concavities. Similarly, results from visual search experiments suggest that local concavities are the crucial features (Hulleman et al., 2000; Humphreys & Müller, 2000). Another example illustrating the importance of concave regions was proposed by Hoffman and Richards (1984). They coined the minima rule, a principle describing how the visual system parses objects into perceptually meaningful parts. The minima rule states that object boundaries are segmented at points of concave indentations (i.e., curvature minima, corresponding to maximum concave curvature) along a contour. These concave regions act as natural part boundaries because they often correspond to physical articulations or joints between object components. In contrast, convex curvature maxima tend to lie within parts rather than marking their separation.

Resnikoff (1985) provided a formal, information-theoretic proof that information is highest in regions of maximum curvature, independent of the sign of curvature. Feldman and Singh (2005) later extended Resnikoff’s proposition and showed that information is highly correlated with curvature. They demonstrated that for closed contours, *concave* regions contain greater information than curvature-matched convex regions (in line with their earlier findings; Barenholtz et al., 2003).^2^ It may be surprising, then, that there is not more evidence of advantages for concave features. However, from the fact that concavities are more informative than convexities when their curvature is matched, it does not follow that *in general* concave regions in a shape will contain more information than convex ones.

### Curvature in neural representation

The psychophysical and theoretical support for a substantial role of curvature information in shape perception align with research on the physiological properties of the primate visual system. Results from electrophysiological studies show that intermediate visual cortical areas in the ventral pathway contain cells tuned to complex polar and hyperbolic gratings and curved stimuli (Gallant et al., 1993; Gallant et al., 1996; Pasupathy, 2006; Pasupathy & Connor, 1999; Pasupathy & Connor, 2001; Pasupathy & Connor, 2002; Yau et al., 2012; Yau et al., 2009). Specifically, the work by Pasupathy and colleagues demonstrated that a large proportion of V4 neurons are sensitive not only to the magnitude and sign of curvature but also to the orientation of the feature (Pasupathy and Connor, 1999, 2001, 2002; see Pasupathy, Popovkina & Kim (2020) for review). Pasupathy and Connor (2002) argued that populations of such curvature-selective V4 neurons could adequately represent whole boundary shapes. Subsequent studies demonstrated that V4 neurons respond to various prominent features, including radial and concentric shapes (David et al., 2006). Carlson et al. (2011) later showed that a population of V4 neurons are curvature-selective with a bias toward acute curvatures. Carlson et al. (2011) suggested that the sparse coding of objects is reflected in the bias of V4 neurons towards less frequent object features, i.e., acute curvature maxima, and a more pronounced representation of uncommon, but salient shape features, which contain most information about object identity. Coding prominent features is efficient from an information-theoretic perspective. Since large portions of natural contours are linear or smoothly curved, while corners and acute curvatures are infrequent, it minimizes redundancy to assign more coding resources to the surprising acute curves than the unsurprising smooth ones (Feldman & Singh, 2005; Timme & Lapish, 2018). More recently, Machida et al. (2024) showed that a significant proportion of primate V4 neurons show tuning for two or more contour features, specifically curvature, closure, symmetry, and orientation, and proposed that V4 neurons do not specialize in just one feature but rather integrate multiple contour attributes simultaneously.

In summary, there is ample physiological and psychophysical evidence that the visual system represents the outline shapes of natural objects on the basis of prominent curvature features. However, there is little consensus about what the critical features are. One theoretical reason to expect a privileged role for extrema is that they tend to carry more information content than low-curvature regions. However, the relationship between curvature extrema and contour information (surprisal) is only correlational. We are not aware of any study that has directly compared how humans make use, in shape tasks, of curvature extrema versus the points of highest surprisal themselves. Most studies assume that the visual system treats some set of curvature features as *special* because they are reliably informative.

Curvature extrema, for example, may be privileged because they are reliable proxies for high-surprisal points on a contour. But an alternative hypothesis is that the visual system does not treat any specific type feature as special *per se*, but rather selectively encodes points that are *informative*. The lack of consensus regarding which curvature features are privileged may reflect a misguided assumption, then. Perhaps what is privileged is information content, which can correlate more or less strongly with different types of curvature features depending on the stimulus domain.

### The current study

This study addresses the question of which curvature properties are the most crucial in shape recognition: convexities, both convexities and concavities, or points of highest information. We tested this by creating pairs of familiar and unfamiliar shapes with varying complexity and testing observers’ ability to match a presented shape with a simplified version thereof. Inspired by Attneave (1954), simplified Attneave-like polygons were created by sampling points along the shape’s boundary and connecting straight lines between those points. We sampled shapes at points of either (i) maximum and minimum curvature, (ii) just maximum curvature, or (iii) highest surprisal. Then, we tested which of these sampling procedures produced simplified polygons that were easiest to match with the original contour. For this, we used two classes of shapes, which we refer to as (i) artificial and (ii) natural shapes. The artificial shapes were composed of combinations of RF patterns (compound RF) of two different levels of complexity. While these patterns can resemble naturally occurring shape outlines such as fruits or human heads (Wilson, Wilkinson, Lin, & Castillo, 2000; Wilson & Wilkinson, 2002; Wilson, Loffler, & Wilkinson, 2002), we previously demonstrated that they can only represent a small subset of planar shapes (Schmidtmann & Fruend, 2019). To overcome these limitations, we also used a variety of animal shapes. Importantly, these also allowed us to disentangle the contribution of points of high curvature and high surprisal, which we achieved by restricting the number of sampled points (see Methods for more information).

In a second experiment, we again sampled shapes at points of either (i) maximum and minimum curvature, or (ii) highest surprisal, however, we restricted the number of sampled points ranging from 4 to 32 points (ranked by signed curvature/surprisal magnitude). If specific contour features are especially important for representing shape, then recognition should still be possible when preserving only a small subset of those features. By constructing simplified polygonal approximations that retain only the most prominent curvature extrema or the most informative (high-surprisal) points, we directly tested which of both carries the most diagnostic information for shape perception.

## Experiment 1

### Methods

#### Observers

Seven observers (mean age: 35, □: ±9.6; 3 females) participated in this study. Three of the authors (FS, NB, GS) were participants in this study. The remaining four were naive as to the purpose of these experiments. GS generated the stimuli, but all target and distractor shapes and their properties were automatically generated and analyzed without the involvement of the author to reduce memory effects. All observers had normal or corrected-to-normal visual acuity. Observations were made under binocular viewing conditions. The experiments were carried out at the University of Plymouth (UK) (N = 2), Loyola University Chicago (USA) (N = 1), and Justus Liebig University Giessen (Germany) (N = 4). Informed consent was obtained from all observers at all institutions. All experiments were approved by the Ethics committees of the above-mentioned institutions and were conducted in accordance with the original Declaration of Helsinki.

#### Apparatus

The generation of the stimuli, the processing of the stimuli and the data analysis were done using MATLAB (R2018a and R2024b, MathWorks, Natick, Massachusetts, USA).

Functions of Psychtoolbox-3 were utilized to present the stimuli (Brainard, 1997; Kleiner et al., 2007). At the University of Plymouth, experiments were run on a Sony Trinitron (Model GDM-F520) CRT monitor with a resolution of 1,280 x 960 and a frame rate of 75 *Hz.* The monitor was controlled by a Mac Mini (2014, 2.8 *GHz*). At Loyola University, Chicago, experiments were run on a Samsung G30A LED Monitor with a resolution of 1,920 x 1,080 and a frame rate of 144 *Hz.* At Justus Liebig University Giessen, experiments were run on an Eizo ColorEdge CG277 LCD Monitor with a resolution of 2,560 x 1,440 and a frame rate of 60 *Hz*. Participants viewed the stimuli at a distance of 40 *cm*. Experiments were performed in dimly illuminated rooms.

### Stimuli

#### Artificial shapes

The artificial shapes were closed contours based on combinations of RF patterns (RF compounds; Wilkinson et al., 1998; Schmidtmann et al., 2019). RF patterns are defined by sinusoidal modulations of a radius in polar coordinates. In general, RF compounds are defined as

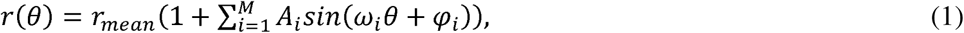

where *r* (radius) and θ refer to the polar coordinates of the contour, and *r_mean_* denotes the radius of the modulated circle and determines the overall size of the contour. *r_mean_* was set to 10, corresponding to a visual angle of ∼4 deg at a viewing distance of 40cm. The variable *A* defines the modulation amplitude (set to 0.2), ω the radial frequency and φ the phase (orientation) for the *i_th_* RF component, respectively. We generated two types of RF compounds, each composed of *M* = 3 RF components (Shape 1: ω*_1_*= 1, ω*_2_* = 2, ω*_3_* = 5; Shape 2: ω*_1_* = 3, ω*_2_* = 5, ω*_3_*= 8). The highest frequency component (ω*_3_*) determines the number of convex maxima and concave minima (i.e., Shape 1 contains 5 and Shape 2 contains 8; see Figure 2). The phase (φ) of each RF component was randomly varied on each trial, such that observers never saw the same shape twice. Thus, observers were presented with a wide range of shapes, varying from shapes with a few, smooth curved boundaries (Shape 1, Figure 2) and shapes with more and relatively salient convexities and concavities (Shape 2, Figure 2). The shapes were black on a white background and had a width of 5 pixels (∼0.2 deg visual angle).

Combinations of different RF patterns have been used previously to describe complex shapes, such as human heads (Loffler et al., 2005; Wilson et al., 2002); however, Schmidtmann and Fruend (2019) demonstrated, experimentally and theoretically, that compound RF patterns represent only a very small and perceptually distinct subset of all possible planar shapes. Due to their mathematical limitations, they are not suited as universal shape descriptors. In order to extend the variety of shapes to more natural shapes, we generated natural closed shapes in the form of animal shapes (which have been demonstrated to be relevant for human behavioral judgements; e.g., Morgenstern et al., 2021).

#### Natural (animal) shapes

The unprocessed natural shapes were based on silhouettes of four animal categories (bear, rabbit, cat, wolf) (Baker & Elder, 2022). The original image files showed black animal silhouettes on a white background (see leftmost column in Figure 1) with varying image dimensions. The original images were resized to 500 x 500 pixels. Most of these original animal silhouettes contain small details, such as fur and whiskers, which introduce high local curvature maxima associated with unimportant features for the overall global shape of the animal. Importantly, the key characteristics of natural shapes can often be captured using just a few of the lowest-frequency components of their Fourier descriptor (FD) representation, allowing for a compact and efficient representation (e.g., Elder, 2018; Wilder, Fruend & Elder, 2018). Therefore, we utilized the FD representation, which is the Fourier transform of the points defining the object boundary, represented as complex numbers (Granlund, 1972), and can be considered as a generalization of the abovementioned RF-based representation that can be applied to arbitrary 2D shapes.

**Figure 1.**
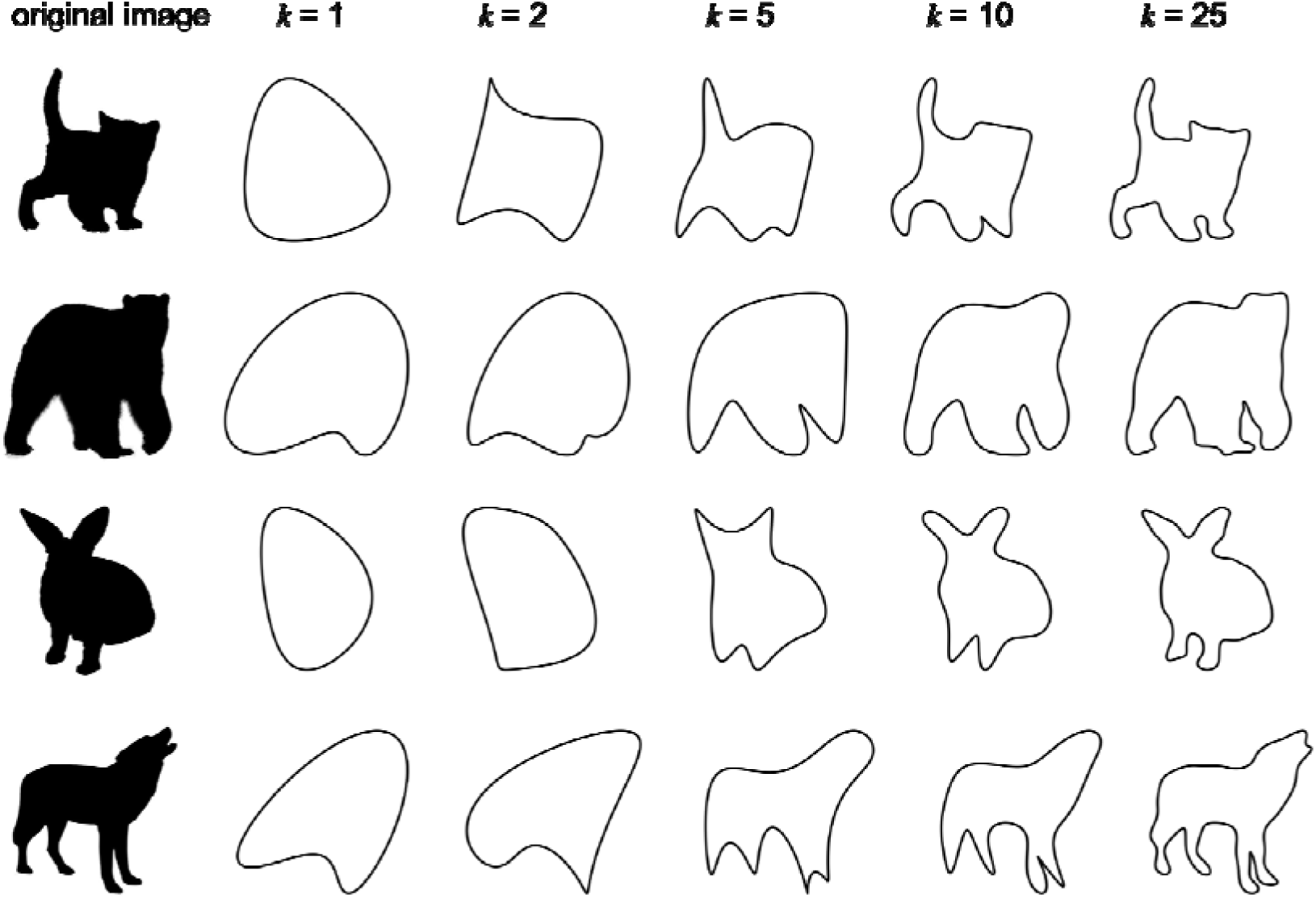
Each row presents an example from the animal categories used in the study. The first column displays the original silhouettes, while the remaining columns show contours using Fourier Descriptors (FDs) with an increasing number of components (k = 1, 2, 5, 10, and 25). The rightmost column shows contours that were used as stimuli in this study, where k = 25.

Figure 1 shows one example of each animal category used in this study. The leftmost column shows the original silhouettes. The contours defined as FDs with an increasing number of Fourier elements (*k* = 1, 2, 5, 10 and 25) are shown in the remaining columns. Briefly, setting k = 25 means that FDs greater than the 26th harmonic will be omitted (see Baker, Wilder & Elder, 2023 for more details). We decided to set k = 25 to satisfy the requirements for this study, i.e., to produce shapes with smooth (un-noisy) outlines that were identifiable as animals. The rightmost column shows contours that were used as stimuli in this study (see further details about the stimulus selection in the Paradigm section below).

### Curvature calculation

The aim of this study was to isolate points of convex and concave curvature maxima along the contours and to connect these points with straight lines. To calculate the curvature along the closed contours, the first and second derivatives were computed using the *gradient* function in MATLAB. The signed curvature□_signed_ along the contour in Cartesian coordinates is defined as

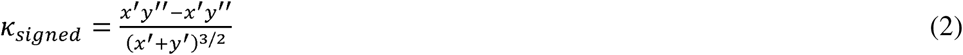

Or

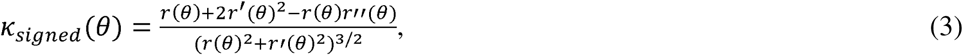

in polar coordinates.

The signed area of the contour (computed with the *Shoelace* formula) was used to determine its orientation. Local convex curvature maxima and concave curvature minima were calculated using MATLAB’s *islocalmax* function. To limit the number of local curvature maxima and minima needed to capture the overall global shape of the animal shapes, we defined a threshold (e.g., K_signed_ > 0.002 for the example shown in Figure 2, bottom). We tested two different curvature conditions: (i) where only the points of local maximum convex curvature (*Convexities-only,* K*_max_*) were connected with straight lines, and (ii) where both the points of maximum convex and maximum concave curvature (K_min_) were connected with straight lines (*Convexities and Concavities*, K_max, min_). Examples of the resulting shapes are illustrated in the second and third columns of Figure 2.

**Figure 2.**
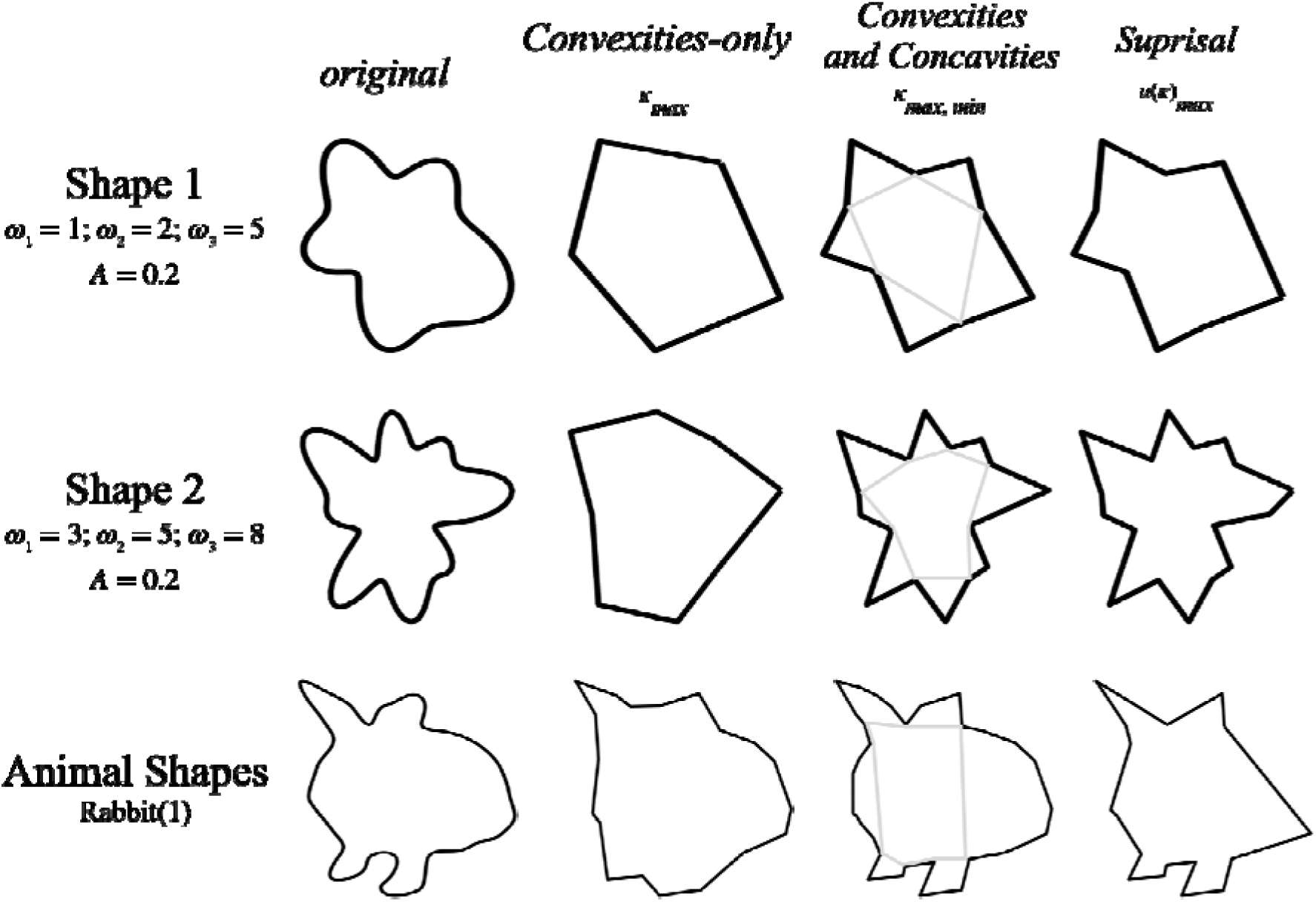
Stimulus examples. The rows show examples for the artificial stimuli based on compound RF patterns with three components with different radial frequencies (Top row, Shape 1: ω1 = 1, ω2 = 2, ω3 = 5; Mid row, Shape 2: ω1 = 3, ω2 = 5, ω3 = 8). The bottom row shows one example of the natural / animal shapes (rabbit). The columns show (1) the original smooth contour (Baseline condition, original), (2) isolated points of maximum convex curvature connected with straight lines (Convexities-only), (3) isolated points of maximum convex and maximum concave curvature connected with straight lines (Convexities and Concavities), and (4) isolated points of maximum surprisal connected with straight lines (Surprisal). Note the differences in the similarity between the Convexities and Concavities and Surprisal conditions for the artificial and natural stimuli. The number of sampled points for the Convexities and Concavities and Surprisal conditions for the Animal

### Maximum surprisal (information)

The rightmost column in Figure 2 shows the condition where points of local information maxima (*Surprisal,* u(K)*_max_*) were connected with straight lines. The surprisal along the closed contour was calculated according to Feldman and Singh’s (2005) method. Given the mathematical definition of the RF compound shapes and their overall symmetrical geometry, the maximum surprisal points often coincide (are highly correlated) with the points of maximum convex and maximum concave curvature (Feldman & Singh, 2005). We calculated the similarity between the stimuli for the *Convexities and Concavities* and *Surprisal* conditions using two common methods, Intersection over Union (IoU, Jaccard index) and the Dice-Sørensen coefficient (Dice, 1945; Sørensen, 1948)^3^, both of which compute similarity as the proportion of overlapping regions between a pair of shapes. This analysis revealed that the similarity between the *Convexities and Concavities* and the *Surprisal* stimuli was higher for the artificial shapes (Shape 1 and Shape 2), compared to the natural shapes (Friedman: ^2^(2) = 4.12, p=.047, IoU) (see examples in Figure 2), which enabled a comparison between the *Convexities-only* and the *Surprisal* condition. To do so, we restricted the number of maximum surprisal points to the number of maximum convex curvature points for each animal shape. Note the differences in the similarity between the *Convexities and Concavities* and *Surprisal* conditions for the artificial and natural stimuli in the examples shown in Figure 2.

### Distractors

The experimental paradigm (see section below for details) required the presentation of a distractor stimulus in each trial. That distractor had to be similar, but sufficiently different from the target stimulus. For the artificial shapes, and in alignment with previous work (Schmidtmann et al., 2015), the distractors were composed of the same RF components as the target shapes, but with different randomly determined phases ( _i_) for each RF component. For the natural shapes, we searched the animal database for the most similar animal shape in each animal category. Similar to the methods described above, the similarity was computed using the Intersection over Union (Jaccard index) and the Dice-Sørensen coefficient (Dice, 1945; Sørensen, 1948). The shape with the highest similarity within the respective animal category was used as the respective distractor shape. The distractor shapes for each condition (*Convexities-only*, *Convexities and Concavities*, *Surprisal*) were generated using the same procedures summarised above (i.e., isolation of points of local curvature maxima and minima or surprisal, and the connection of these points with straight lines). Examples of distractor shapes are shown in Figure 3.

**Figure 3.**
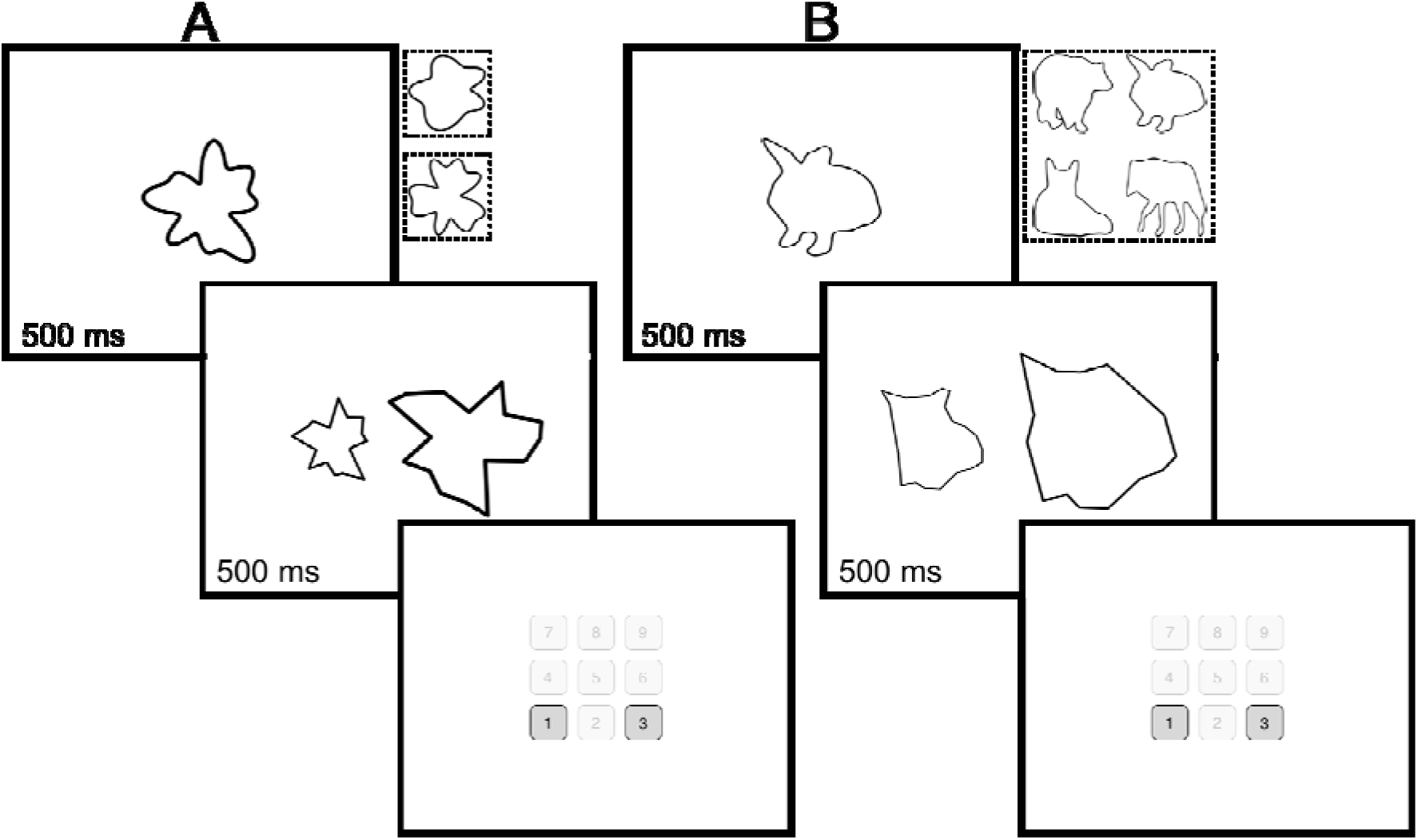
The figure shows the experimental Match-to-Sample paradigm for (A) artificial and (B) natural shapes. The task in an experimental trial was to match a smooth shape (test) to two subsequently presented shapes (scaled in size), the target and distractor. The target shape was composed by isolating (i) points of maximum convex curvature, (ii) points of maximum convex and maximum concave curvature, and (iii) points of maximum surprisal, connected with straight lines. The position (left or right) of the target and distractor shape was randomly assigned in each trial. Conditions (i) – (iii) were randomly presented in each experimental block. The artificial and natural shapes were tested in separate blocks. Two artificial shape classes and four different natural (animal) shape categories were tested randomly in each experimental block. Examples of these shape categories are shown in the additional icons next to the top panel in (A) and (B), where the dashed squares indicate the blocks. The task for the observer was to decide if the target shape was presented on the left or right by pressing the keys 1 (left) or 3 (right) on a numeric keypad.

All stimuli and the experimental code are openly accessible here.

### Paradigm

Using a Match-to-Sample paradigm, the task for the observers was to match a smooth shape (test) to one of two subsequently presented shapes, the target and distractor. An experimental trial was initiated by the presentation of the test shape in the center of the screen for 500 *ms*, which was immediately followed by the simultaneous side-by-side presentation of the distractor and target shapes (500 *ms*). The target and distractor shapes were each randomly scaled in size by ±1/3 of the test shape, and their locations were randomly determined on a trial-by-trial basis (left or right side). The scaling was applied to minimize the possibility of observers simply judging the distance between the isolated points of curvature maxima, minima and maximum surprisal. There is good evidence that this scaling does not significantly impair performance due to the scale-invariant nature of shape recognition (e.g., Schmidtmann et al., 2016; Wilkinson et al., 1998).

The position (left or right) of the target and distractor shape was randomly set in each trial. All conditions (*Baseline*, *Convexities-only*, *Convexities and Concavities*, *Surprisal*) were randomly interleaved in an experimental block. The artificial and natural shapes were tested in separate blocks. The two classes of artificial shapes (Shape 1, Shape 2) were tested in separate blocks, each subdivided into four smaller blocks of 50 presentations to limit test duration, so that each observer was presented with an overall number of 400 different shapes (200 for each artificial shape category). This high number of different shapes was possible because varying the phase of the *i_th_* RF component changes the overall shape of the resulting compound shape. However, the number of natural shapes for each animal category in the database was limited. Given the characteristics of the original animal shapes in the database (e.g., image quality, clearly identifiable animal category, non-overlapping features (e.g., legs, head, tail), animal body posture, etc.), we selected ten images from four different animal categories (bear, rabbit, cat, wolf). Each of these images was randomly tested 100 times, leading to an overall number of 400 trials. These trials were subdivided into two smaller blocks (200 presentations) to reduce testing time. Examples of these shape categories are shown in the additional icons next to the top panel in Figure 3A and B. The red squares indicate the experimental blocks. The presentation of the target and distractor shapes was followed by a blank white screen, during which observers were asked to choose whether the target shape was presented on the left or the right, by pressing the keys 1 (left) or 3 (right) on a numeric keypad. Observers were instructed to respond as quickly as possible. The response time was recorded for subsequent analysis. The key press initiated the next experimental trial. No feedback was provided for incorrect responses. The experimental paradigm is illustrated in Figure 3.

### Results

#### Artificial shapes

Figure 4 shows the results for the artificial shapes. Qualitatively, the overall pattern is similar for both shape classes (Figure 4 left vs. right). On average, the lowest performance was measured for the condition where only points of convex maxima were connected with straight lines (*Convexities-only*; Shape 1: 67 ±13%; Shape 2: 65 ±12%), followed by the baseline condition, where the test shape and target shape were identical (*Baseline*; Shape 1: 84 ±14%; Shape 2: 88% ±8%). Finally, the best and similar performances were measured for the conditions with both convex maxima and concave curvature minima (*Convexities and Concavities;* Shape 1: 87 ±10%, Shape 2: 89 ±6%) and the condition where points of high surprisal were isolated and connected with straight lines (*Surprisal*; Shape 1: 85 ±8%, Shape 2: 93 ±4%).

**Figure 4.**
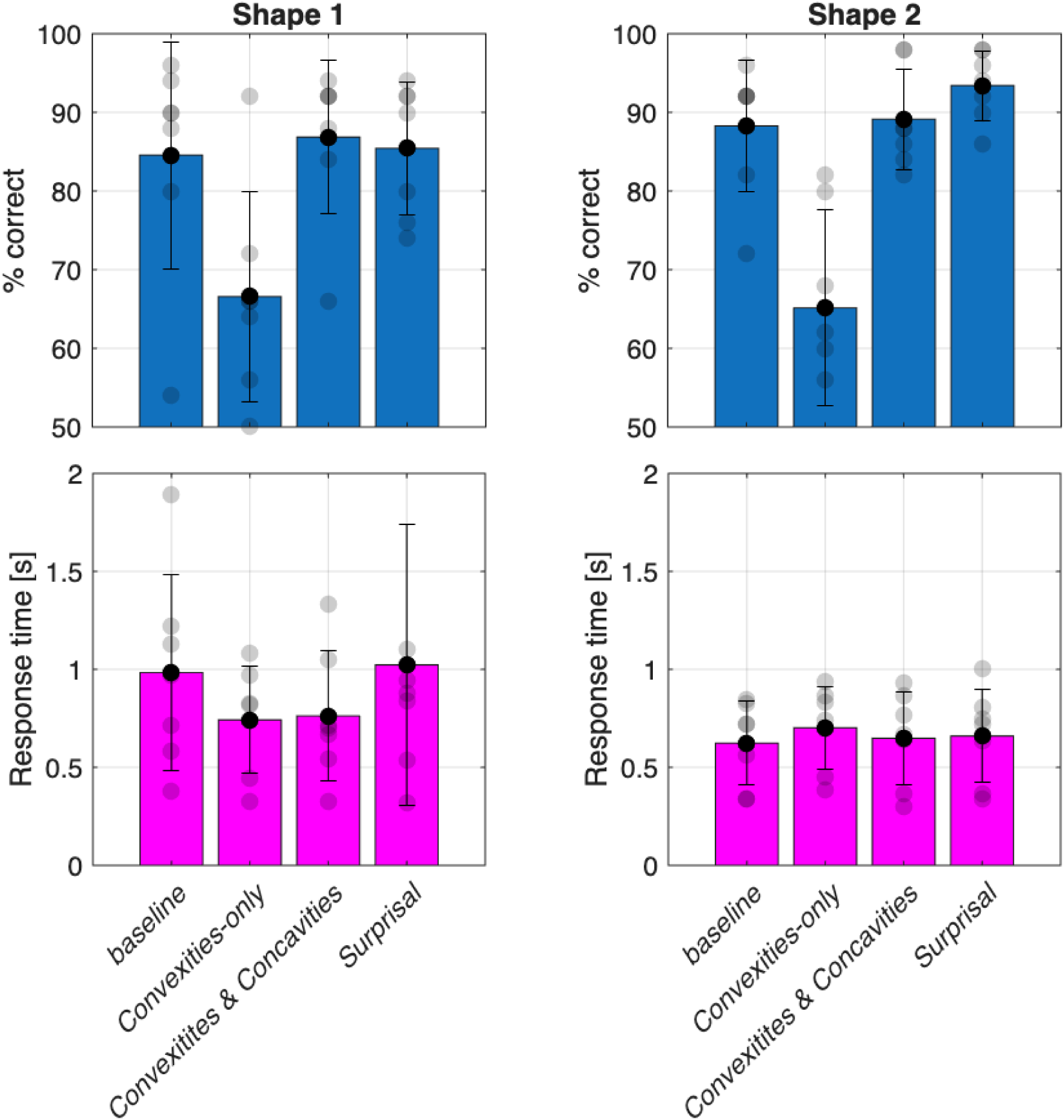
The top row shows performance (% correct) and the bottom row the response time for Shape 1 (left) and Shape 2 (right) for each condition (Baseline, Convexities-only; Convexities and Concavities; Surprisal). The bottom row shows the corresponding mean and individual response times [s]. The height of the bars and the solid black data points show the mean results across observers, the error bars the standard deviations, and the light grey data points the individual results.

Note that the performance in the *Convexities and Concavities* and the *Surprisal* conditions is statistically not significantly different from the baseline, which is surprising given that in the latter the test shape was identical to the target shape. If anything, the performance in the *Convexities and Concavities* and the *Surprisal* condition are descriptively slightly higher than baseline.

#### Natural shapes

Figure 5 shows the results for the natural shapes. The overall pattern is very similar to the results for the artificial shapes: Here, descriptively, the best average performance was measured for the baseline condition (*Baseline*, 95 ±3%), followed by a similar performance for *Surprisal* (90 ±4%) and the *Convexities and Concavities* condition (87 ±7%). As for the artificial shapes, the worst performance was obtained for the *Convexities-only* condition (67% ±7%). A particularly interesting result is the similar performance for the *Convexities and Concavities* and *Surprisal* conditions (87% vs. 90%), despite the lower number of points in the *Surprisal* condition (as a result of equating the points in the *Surprisal* and *Convexities-only* condition).

**Figure 5.**
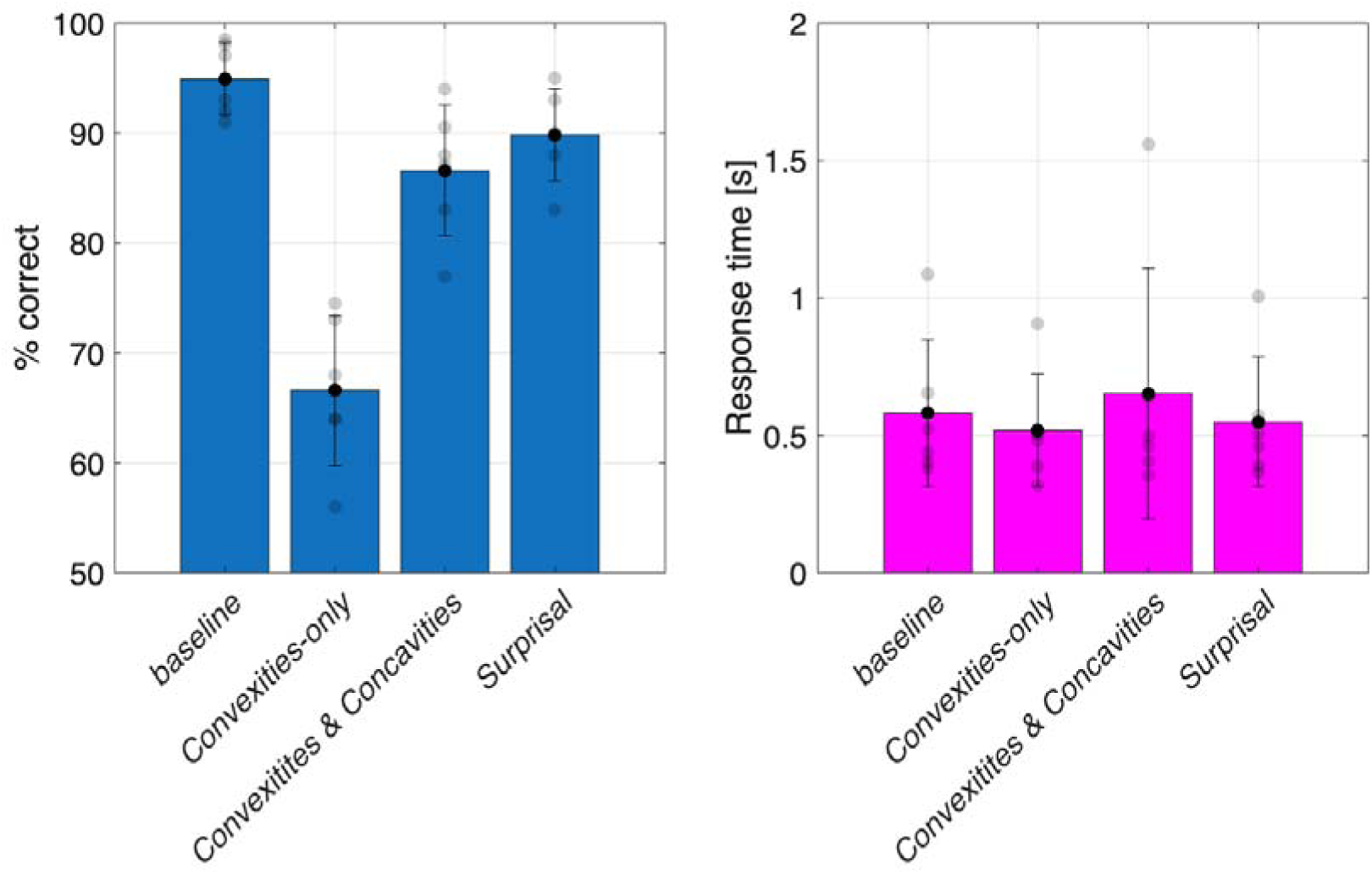
The left graph shows performance (% correct) for the natural shapes for each condition (Baseline, Convexities-only; Convexities and Concavities; Surprisal). The right graph shows the mean and individual response times [s]. The height of the bars and the solid black data points show the mean results across observers, the error bars the standard deviations, and the light grey data points the individual results.

Results for artificial and natural shapes were statistically analyzed using a non-parametric Friedman ANOVA with subsequent pairwise comparisons (Bonferroni corrected). This analysis revealed a statistically significant global effect (X^2^(11) = 46.4, p<.001). Statistically significant pairwise comparisons are summarized in Table A1. The key results are that in all conditions, (i) the performance for the *Convexities-only* condition was significantly poorer compared to the baseline, (ii) there was in almost all cases no statistically significant difference between the *Baseline* and the *Convexities and Concavities*, and *Surprisal* conditions (2 significant out of 6 total comparisons). (iii) Overall, there is no significant difference between the artificial and natural shapes.

#### Response times

The bottom panel in Figure 4 and the right panel in Figure 5 show the corresponding mean and individual response times [s]. The height of the bars and the solid black data points show the mean results across observers, the error bars the standard deviations, and the light grey data points the individual results. Results show that response times for Shape 1 (range: 0.74 ±0.3s to 1.02 ±0.7s) were, on average, statistically significantly higher compared to Shape 2 (range: 0.62 ±0.2s to 0.7 ±0.2s) and the natural shapes (range: 0.51 ±0.2s to 0.65 ±0.4s) (see Table A2 for details). In other words, increasing the complexity, i.e., the number of curvature maxima and minima, seems to increase the speed of decision-making, but not necessarily performance. However, more importantly, the response times did not significantly differ between the *Baseline*, *Convexities-only*, *Convexities and Concavities*, *Surprisal* conditions.

#### Inverse Efficiency Score

For each observer and condition (*Baseline*, *Convexities-only*, *Convexities and Concavities*, *Surprisal*), inverse efficiency scores (IES) were computed by dividing mean response time of correct responses by the proportion correct, thereby combining response time and accuracy into a single performance measure (Figure 6). The IES were analyzed separately for each stimulus class using repeated-measures ANOVAs. Results showed no significant effect of sampling method for Shape 1, *F*(3,18) = 1.54, *p* = .261, or for the animal shapes, *F*(3,15) = 2.43, *p* = .177, but for Shape 2, *F*(3,15) = 16.44, *p* < .001(Greenhouse–Geisser corrected). Subsequent pairwise comparisons indicated that for Shape 2 the IES was significantly higher in the *Convexities-only* compared to the *Baseline* (*p* = .005), the *Convexities and Concavities* (*p* = .017), and the *Surprisal* (*p* = .019) conditions, whereas *Baseline*, *Convexities and Concavities*, and *Surprisal* conditions did not differ significantly from one another. In conclusion, the performance efficiency was reduced in the *Convexities-only* for Shape 2, but not affected by condition for Shape 1 or with the animal stimuli. This replicated our results in the response times, suggesting that those were not explained by speed-accuracy trade-offs.

**Figure 6.**
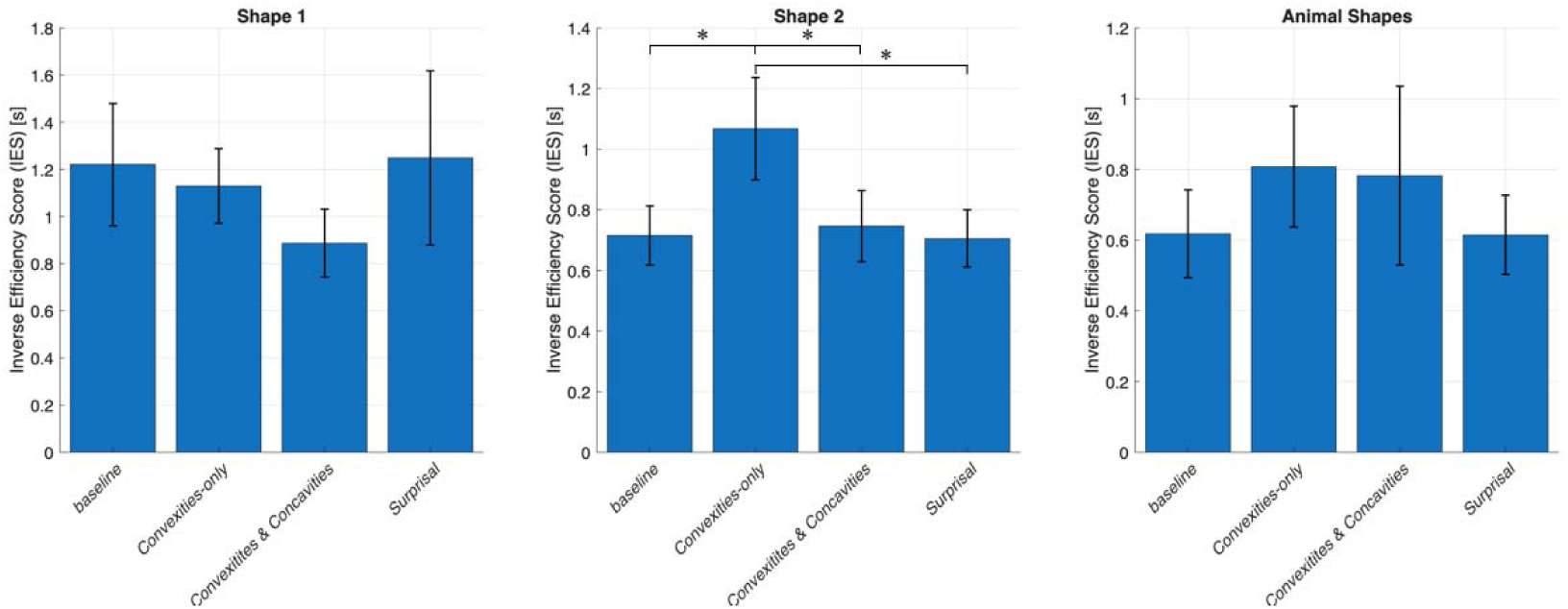
Inverse efficiency scores (IES), computed by dividing the mean reaction time by proportion correct, thereby combining response time and accuracy into a single performance measure. For other details see Figures 4 and 5.

### Experiment 2

#### Stimuli

Stimuli were identical to the animal shapes used in Experiment 1 (see Natural (animal) shapes section above). Briefly, local contour curvature was computed according to eq. 2. Points of maximum convex and concave curvature were identified as local extrema (*Convexities and Concavities*) and surprisal was computed along the contour to identify points of maximal informational content (*Surprisal*). For each stimulus, these points were rank-ordered based on either the magnitude of signed curvature or surprisal, and the highest *n* points were selected. We tested seven conditions with points of *n* = 4, 6, 8, 12, 16, 24 and 32. To preserve the geometric structure of the contour, the selected points were re-ordered according to their sequential position along the contour and connected by straight lines, resulting in simplified polygonal approximations of the original shape. Example stimuli are shown for one cat shape in Figure 7.

**Figure 7.**
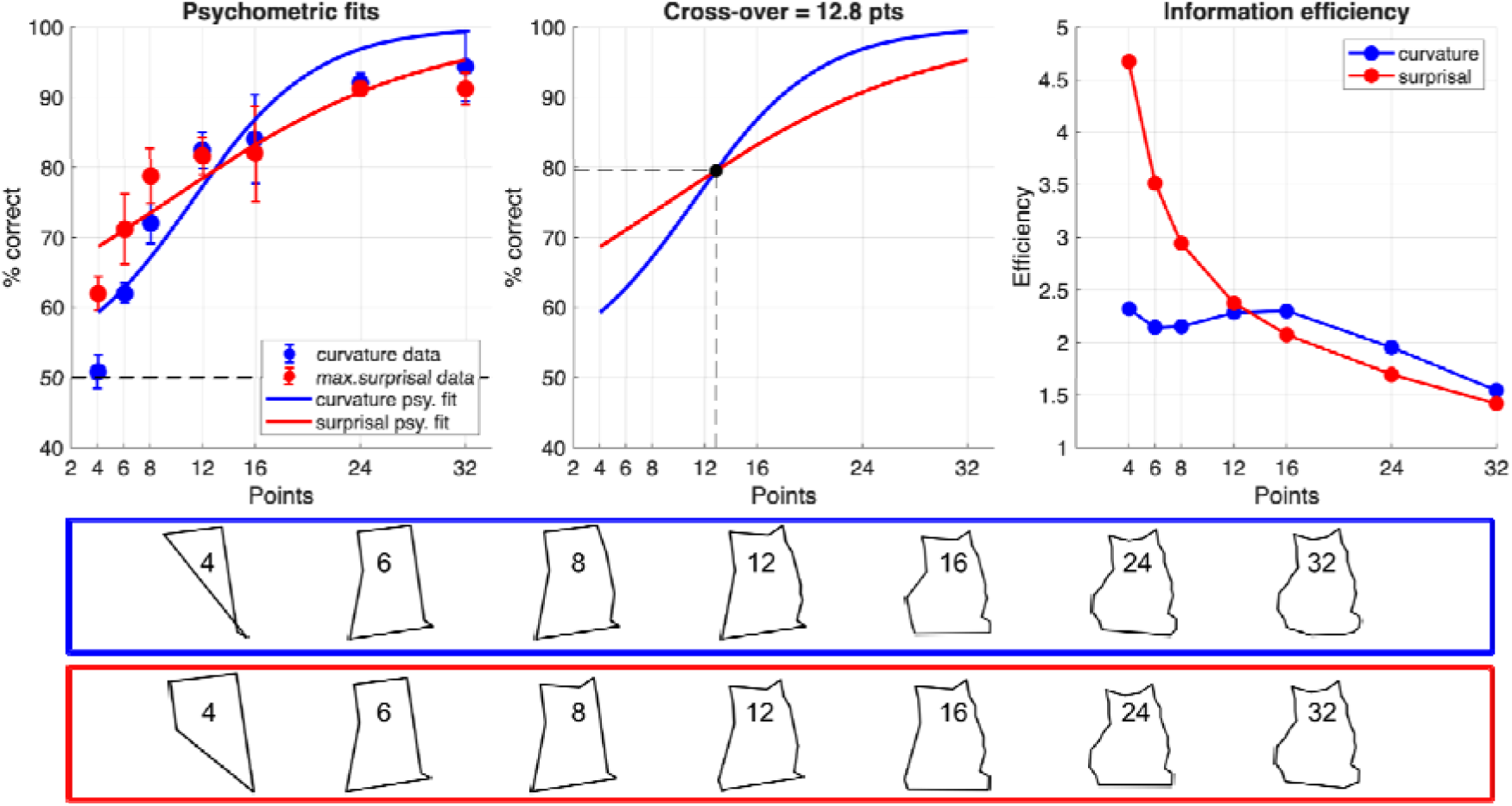
The graph shows the results for Experiment 2. The leftmost graph shows the average performance (% correct) as a function of the number of sampled points (n = 4 - 32) across five observers. The blue data points show results for the Convexities and Concavities condition (curvature data), and the red data points for the Surprisal condition (max. surprisal data). The error bars represent the standard deviation, and the dashed line shows chance performance (50%). Psychometric functions (Logistic, left) were fit to the data, and the cross-over point was calculated (middle, dashed line; see text for details). The rightmost graph shows information efficiency for curvature- and surprisal-based sampling. The icons below the graph show stimulus examples for the Convexities and Concavities condition in the blue box and the Surprisal condition in the red box. The number within each shape refers to the number of sampled points.

#### Paradigm

Similar to Experiment 1, we used a match-to-sample paradigm, where five observers (authors GS, NB, FS and two naive observers) were matching a smooth reference animal shape to one of two subsequently presented rescaled shapes (target and distractor), that were constructed by either extracting the points of (1) maximum and minimum curvature (*Convexities and Concavities*), or (2) maximum information (*Surprisal*) and connect them with straight lines. All conditions were randomly interleaved.

### Results

The data points in the leftmost graph in Figure 7 show the average performance across observers for the *Convexities and Concavities* condition in blue and the *Surprisal* condition in red. The error bars show standard deviations. Performance increased with increasing number of points in both conditions, ranging on average from an initial 50% for the *Convexities and Concavities* condition and 60% for the *Surprisal* condition for shape composed of 4 sampled points to a performance of ∼90% in both conditions for 24 sampled points. Results were analyzed using a linear mixed-effects model on logit-transformed proportions, with fixed effects of condition, levels (number of points *n*), and their interaction, and a random intercept for subject. The model revealed a significant effect of level, reflecting an increasing performance with an increasing number of points, and a condition × level interaction, indicating that differences between conditions varied across levels. Paired t-tests showed that for the lowest two levels (*n*=4, 6), the *Surprisal* condition (62%, 71%) significantly outperformed the *Convexities and Concavities* condition (51%, 62%), (*p* = .027, *p* = .034; Holm-corrected). No reliable statistically significant between-condition differences were observed at higher levels (all corrected *p* > .05), indicating veridical performance. One-sample t-tests against chance (50%) showed that the performance for the *Surprisal* condition exceeded chance at the lowest level, *t*(4) = 8.47, *p* < .001, whereas the *Convexities and Concavities* condition did not, *t*(4) = 0.76, *p* = .48. Both conditions were significantly above chance at all subsequent levels (all *p* < .05, corrected). These results demonstrate an early performance advantage for the *Surprisal* condition.

The data were fitted with psychometric functions (Logistic) using the Palamedes Toolbox (Prins & Kingdom, 2018) with fixed asymptotes. The threshold and slope parameters were assessed using a non-parametric bootstrap procedure. Subjects were resampled with replacement (5000 iterations), and psychometric functions were refitted to the resampled data on each iteration. Distributions of parameter differences (curvature minus surprisal) were used to derive 95% confidence intervals (CI) and two-tailed *p*-values. Results show a clear dissociation between sampling strategies. Surprisal-based sampling exhibited a significantly lower threshold than curvature-based sampling (Δ = 1.91, 95% CI [0.57, 3.96], *p* < .001), indicating more efficient performance at low sampling densities. Curvature-based sampling showed a significantly steeper slope (Δ = 0.109, 95% CI [0.079, 0.165], *p* < .001), reflecting a stronger improvement in performance as the number of points increased. This trade-off resulted in a cross-over point at approximately 12.8 sampled points (∼79.5% correct; dashed line in the middle panel of Figure 7), marking the transition from an initial advantage of surprisal-based sampling to superior performance for curvature-based representations at higher sampling densities. Consistent with this pattern, an information-efficiency analysis, (*performance* − *chance*)/*points*, showed that surprisal-based sampling provided substantially greater efficiency at fewer points (i.e., 4–8 points, rightmost graph in Figure 7), whereas curvature-based sampling became more efficient at higher densities (≥12 points). Together, these results suggest that information-theoretic sampling supports efficient initial shape encoding under sparse conditions, while curvature-based representations scale more effectively as contour resolution increases.

## Discussion

There is a significant body of evidence suggesting that prominent boundary features, such as points of high curvature, are the basis for the representation of natural objects in human vision. Given the partly conflicting evidence about the importance of either convex/concave curvature features or points of high information along the contour, we conducted shape-matching experiments, where observers matched a smooth Attneave-like test shape (artificial or animal shape) to one of two subsequently presented re-scaled shapes (target and distractor). The target and distractor shapes were created by isolating the points of (i) maximum curvature, (ii) maximum and minimum curvature, or (iii) maximum information, and connecting them with straight lines.

### Importance of shape complexity and familiarity

Results from Experiment 1 show that overall performance is similar for the two classes of artificial shapes, despite their differences in complexity. Shape 1 contained RF components with the radial frequencies of ω*_1_* = 1, ω*_2_* = 2, ω*_3_* = 5, and Shape 2 frequencies of ω*_1_* = 3, ω*_2_* = 5, ω*_3_* = 8. The highest frequency in each RF compound (i.e., ω*_3_*) determines the number of curvature maxima and minima. Hence, Shape 1 contained five, and Shape 2 eight convex maxima and concave minima (see Figure 2). Given the difference in complexity (i.e., number of perturbations), one might expect performance and response times to depend on the number of curvature maxima; however, the results do not show this dependency, neither for the *Baseline* condition, nor the *Convexities and Concavities*, and *Surprisal* conditions. While previous studies on RF discrimination, that is the ability to discriminate between a circle and a slightly modulated RF pattern, showed that thresholds are dependent on the radial frequency, where thresholds decrease with increasing radial frequency and curvature maxima (Wilkinson et al., 1998; Schmidtmann & Kingdom, 2017), our results show that this is not the case for the shape recognition task used here.

Given their mathematical definition, i.e., the sinusoidal modulation of a radius from a common center point, compound RF patterns, especially with low amplitudes used here, are largely circular, with only subtle salient protrusions and are often symmetrical (depending on their individual phases; Schmidtmann, Kingdom & Loffler, 2019). Previous work also showed that these RF compound shapes can only represent a small subset of all planar shapes (Schmidtmann & Fruend, 2019). We aimed to investigate how the results differ between artificial shapes and shapes that resemble more natural, animal forms, which do not have the aforementioned limiting characteristics of RF-based shapes. The natural shapes used here are more complex than the RF-based shapes. They are asymmetrical and contain prominent (protruding) features, such as long, thin legs, long tails, rabbit ears, etc., typical of the animals used here. Further, recall that the most informative regions in natural shapes are not entirely restricted to concave regions. Surprisingly, while performance is slightly higher for natural compared to artificial shapes, the difference is relatively small, and the overall pattern of results across all conditions is very similar. Hence, our data suggest that observers’ familiarity with the natural shapes does not significantly improve performance. This is interesting, given the previously reported advantage of familiarity in the recognition of faces (Hancock et al., 2000; Anaki & Bentin, 2009), objects (Noudoost et al., 2005), and visual search, where search was faster for familiar stimuli (Wang, Cavanagh & Green, 1994; Greene & Rayner, 2001; but see Madrid et al., 2019).

### Importance of convexities and concavities and comparison to earlier findings

Another interesting observation is that across all shapes and conditions, observers’ performance was significantly lower for the condition where the points of maximum convex curvature were connected with straight lines (*Convexities-only*), compared to the *Baseline* condition and the conditions where either both, convex maxima and concave minima (*Convexities and Concavities*) or the points of maximum surprisal (*Surprisal*) were connected with straight lines. Lower performance for shapes composed of points of maximum convex curvature contradicts previous findings of a superior performance for convexities in shape recognition (see introduction for review). Also, the results reported here challenge Schmidtmann et al.’s (2015) previous observations, even though stimuli and experimental paradigm are similar. However, while their stimuli were equivalent to the RF-based artificial shapes used here (they did not use natural shapes), observers were presented with a segmented test shape (reference shape), which only showed either points/regions of maximum convexities, concavities or intermediate regions of varying segment lengths (ranging from single points to longer segments). The regions between these points were omitted rather than connected with straight lines as our stimuli. The task was to match this segmented shape to one of two subsequently presented whole-contour shapes (target and distractor). Their results showed that for very short segment lengths (single points), performance was significantly higher for convexities (∼80%) compared to concavities or intermediate regions (∼60%) and was independent of segment length. For concavities and intermediate points, performance for very short segments was at chance level and improved with increasing segment length, reaching convexity performance only for long segments. Schmidtmann et al. (2015) concluded that for this shape-matching experiment, closed curvilinear shapes are encoded using the positions of convexities, rather than concavities or intermediate regions. The superior and segment-length-independent performance of convex maxima and the unique characteristics of shapes that result from connecting these points inspired Schmidtmann et al. (2015) to develop a model that provided a very good account of their data.^4^ Briefly, their model proposes that during an experimental shape-matching trial, the visual system employs a rudimentary template shape constructed by connecting the endpoints of the convex, concave or intermediate segments in the target and distractor shape with straight lines. This template shape is then compared to the whole-contour target shape and the distractor shape. The decision rule is to select the shape (target or distractor) most similar to the test shape template. This model does not generalize to designs where the observer is not initially presented with the isolated curvature features. In the current experimental paradigm, on the other hand, the observers were presented with the smooth whole-contour shape; hence, we would not expect the visual system to create a template shape connecting specific curvature features with straight lines. However, the idea of a perceptual similarity comparison between the smooth test shape and the target and distractor shapes would still be possible.

Another interesting observation is that the performance for the *Convexities-only* condition in the current study is considerably lower than that of the comparable shortest segment length condition in Schmidtmann et al.’s study (∼60% vs. ∼80%). This is counterintuitive given that the stimuli here contained more visible shape information in the form of straight-line connections between the points of maximum convex curvature, whereas Schmidtmann et al.’s stimuli omitted these regions. The suggestion that the addition of these visible straight lines impairs performance seems unlikely. One might speculate that the abovementioned differences in the experimental paradigm could be responsible. As discussed earlier, the importance of convex, concave or intermediate regions depends on stimulus type and experimental paradigms.

We can also show that, on average, observers’ results are similar for the *Convexities and Concavities*, and *Surprisal* compared to the *Baseline* condition across all shapes. This suggests that crucial information can be extracted quickly (within 500ms) from simplified shapes that only contain points of maximum convex and concave curvature, despite all the missing intermediate shape information. In line with previous studies that show that Attneave-like shapes constructed by connecting points of high curvature are easier to identify compared to polygons created by connecting intermediate points or self-identified salient points (De Winter & Wagemans, 2008a), the results here support Attneave’s (1954) original hypothesis about the importance of maximum curvature points in shape representation.

The results presented in the current study reveal a noteworthy advantage for high-surprisal points. The similar performances for the *Convexities and Concavities* and the *Surprisal* conditions observed in Experiment 1 are understandable for the artificial shapes, given that surprisal is very closely correlated with curvature for these shapes (Feldman & Singh, 2005). However, these results are less intuitive for the natural shapes, where we were able to disentangle curvature from surprisal. Remember that for the natural shapes, we equated the number of maximum surprisal points to the number of maximum convex curvature points. Yet, performance in the *Convexities and Concavities* and the *Surprisal* conditions are similar. In other words, for the natural shapes, fewer points of high surprisal (*Surprisal*; ∼60%)^5^ are required to achieve the same performance as with points of maximum convex and concave (*Convexities and Concavities*), suggesting that at least for the natural shape stimuli used here, the points along the contour that contain high information are relatively more important.

In Experiment 2, we measured matching performance for even more simplified polygonal approximations of the original shape by systematically varying the number of sampled points (4–32). Results show that performance generally increased with number of points in both conditions (Convexities and Concavities & Surprisal). However, simplified shapes composed of points of high surprisal exhibited significantly lower thresholds than shapes composed of points of high curvature – indicating superior performance at low sampling densities. At the same time, shapes based on curvature sampling showed a significantly steeper slope of the psychometric functions, reflecting a stronger improvement in performance as the number of points increased. We estimate the transition from the initial advantage of surprisal-based sampling to that of curvature-based sampling at about 13 sampled points. Additional information-efficiency analysis showed that surprisal-based sampling provided substantially greater efficiency at fewer points (i.e., 4–8 points), whereas curvature-based sampling became more efficient at higher densities (≥12 points). These results indicate that early representations prioritise informationally diagnostic features, while later stages increasingly rely on structured geometric descriptions of object boundaries.

Overall, the results from these experiments suggest that the visual system might use regions of higher information rather than maximum curvature itself when representing shapes.

### Limitations

It should be noted that the assumptions underlying the present approach are restricted to smooth, two-dimensional planar shapes defined by a one-dimensional boundary. While such representations offer experimental tractability, they necessarily oversimplify real-world object perception. In natural vision, objects are frequently partially occluded, self-occluding, or embedded in complex scenes, such that shape information cannot be recovered from a complete outline alone. Consequently, models based purely on boundary extraction provide only a limited account of shape representation. Additional visual cues, including shading, perspective, texture gradients, and depth information, also play a critical role in object recognition and must be incorporated into more comprehensive encoding frameworks.

Developing such integrative models remains an important goal for future research.

### Conclusion and outlook

How do our results relate to existing theories of shape recognition and representation? As mentioned in the introduction, there is considerable psychophysical and physiological evidence that shapes are encoded by a surface-based, object-centered representation in the intermediate-level processing in a hierarchical computation (Kubilius et al., 2014). Neurons along the ventral stream show tuning to curvatures, orientations, and contours. Contours, i.e. the outlines of object shapes, provide crucial information for the construction of the object-based representation (Attneave, 1954; Elder et al., 2018). Psychophysical studies have reported the crucial role of complex contour features, including closure, convexity, and symmetry, in the representation (Bertamini and Wagemans, 2013). The superior and shape-independent performance for convex shape features observed here is supportive of Carlson et al.’s (2011) hypothesis that shapes might be encoded from specific salient shape features. Carlson et al. and previous work (see Pasupathy, Popovkina & Kim (2020) for review) demonstrated that V4 neurons are tuned to high convexities and concavities rather than to shallow convex or concave curves. Carlson et al. (2011) further demonstrated that common objects mainly contain smooth curvature (acute curvatures are rare), while V4 neurons are tuned to acute curvature (convexities and concavities). They suggested that efficient/sparse coding of general shapes is achieved by biasing representation towards features with high information content but a lower probability of occurrence. This encoding scheme of object boundaries first emerges in V4 and subsequently in V1, possibly via feedback (Chen et al., 2014). Our results provide further psychophysical evidence for this shape encoding scheme in terms of high curvature points, with additional evidence that points of high information are of specific importance. Given that both features seem to diverge to some extent in natural shapes, future studies might further aim to disentangle their roles for our representation of shapes.

## Appendix

**Table A1.** The table shows the statistically significant pairwise comparisons in performance (*p*<.05; Friedman ANOVA, Bonferroni corrected). K_max_: *Convexities-only*, K_max,min_: *Convexities and Concavities*, u(K)_max_: *Surprisal*

| Pairwise Comparisons |  | Statistic | p-value |
| --- | --- | --- | --- |
| Shape 1 *baseline* | Shape 1 $\kappa_{\max}$ | 3.9983 | $< \square .001$ |
| Shape 1 *baseline* | Shape 2 $\kappa_{\max}$ | 4.6082 | $< \square .001$ |
| Shape 1 *baseline* | Natural shape *baseline* | 2.8463 | 0.006 |
| Shape 1 *baseline* | Natural shape $\kappa_{\max}$ | 4.676 | $< \square .001$ |
| Shape 1 $\kappa_{\max}$ | Shape 1 $\kappa_{\max, \min}$ | 4.4049 | $< \square .001$ |
| Shape 1 $\kappa_{\max}$ | Shape 1 $u(\kappa)_{\max}$ | 3.1173 | 0.003 |
| Shape 1 $\kappa_{\max}$ | Shape 2 *baseline* | 4.4049 | $< \square .001$ |
| Shape 1 $\kappa_{\max}$ | Shape 2 $\kappa_{\max, \min}$ | 4.4049 | $< \square .001$ |
| Shape 1 $\kappa_{\max}$ | Shape 2 $u(\kappa)_{\max}$ | 5.557 | $< \square .001$ |
| Shape 1 $\kappa_{\max}$ | Natural shape *baseline* | 6.8446 | $< \square .001$ |
| Shape 1 $\kappa_{\max}$ | Natural shape $\kappa_{\max, \min}$ | 2.1008 | 0.04 |
| Shape 1 $\kappa_{\max}$ | Natural shape $u(\kappa)_{\max}$ | 3.8628 | $< \square .001$ |
| Shape 1 $\kappa_{\max, \min}$ | Shape 2 $\kappa_{\max}$ | 5.0148 | $< \square .001$ |
| Shape 1 $\kappa_{\max, \min}$ | Natural shape $\kappa_{\max}$ | 5.0826 | $< \square .001$ |
| Shape 1 $u(\kappa)_{\max}$ | Shape 2 $\kappa_{\max}$ | 3.7272 | $< \square .001$ |
| Shape 1 $u(\kappa)_{\max}$ | Shape 2 $\kappa_{\max}$ | 2.4397 | 0.018 |
| Shape 1 $u(\kappa)_{\max}$ | Natural shape *baseline* | 3.7272 | $< \square .001$ |
| Shape 1 $u(\kappa)_{\max}$ | Natural shape $\kappa_{\max}$ | 3.795 | $< \square .001$ |
| Shape 2 *baseline* | Shape 2 $\kappa_{\max}$ | 5.0148 | $< \square .001$ |
| Shape 2 *baseline* | Natural shape *baseline* | 2.4397 | 0.018 |
| Shape 2 *baseline* | Natural shape $\kappa_{\max}$ | 5.0826 | $< \square .001$ |
| Shape 2 *baseline* | Natural shape $\kappa_{\max, \min}$ | 2.3041 | 0.025 |
| Shape 2 $\kappa_{\max}$ | Shape 2 $\kappa_{\max}, \kappa_{\min}$ | 5.0148 | $< \square .001$ |
| Shape 2 $\kappa_{\max}$ | Shape 2 $u(\kappa)_{\max}$ | 6.1669 | $< \square .001$ |
| Shape 2 $\kappa_{\max}$ | Natural shape *baseline* | 7.4545 | $< \square .001$ |
| Shape 2 $\kappa_{\max}$ | Natural shape $\kappa_{\max, \min}$ | 2.7107 | 0.009 |
| Shape 2 $\kappa_{\max}$ | Natural shape $u(\kappa)_{\max}$ | 4.4727 | $< \square .001$ |
| Shape 2 $\kappa_{\max, \min}$ | Natural shape *baseline* | 2.4397 | 0.018 |
| Shape 2 $\kappa_{\max, \min}$ | Natural shape $\kappa_{\max}$ | 5.0826 | $< \square .001$ |
| Shape 2 $\kappa_{\max, \min}$ | Natural shape $\kappa_{\max, \min}$ | 2.3041 | 0.025 |
| Shape 2 $u(\kappa)_{\max}$ | Natural shape $\kappa_{\max}$ | 6.2347 | $< \square .001$ |
| Shape 2 $u(\kappa)_{\max}$ | Natural shape $\kappa_{\max, \min}$ | 3.4562 | 0.001 |
| Natural shape *baseline* | Natural shape $\kappa_{\max}$ | 7.5223 | $< \square .001$ |
| Natural shape *baseline* | Natural shape $\kappa_{\max, \min}$ | 4.7438 | $< \square .001$ |
| Natural shape *baseline* | Natural shape $u(\kappa)_{\max}$ | 2.9818 | 0.004 |
| Natural shape $\kappa_{\max}$ | Natural shape $\kappa_{\max, \min}$ | 2.7785 | 0.007 |
| Natural shape $\kappa_{\max}$ | Natural shape $u(\kappa)_{\max}$ | 4.5405 | $< \square .001$ |

**Table A2.** The table shows the statistically significant pairwise comparisons in response times (*p*<.05; Friedman ANOVA, Bonferroni corrected). K_max_: *Convexities-only*, K_max,min_: *Convexities and Concavities*, u(K)_max_: *Surprisal*

| Pairwise Comparisons |  | Statistic | p-value |
| --- | --- | --- | --- |
| Shape 1 *baseline* | Shape 2 $\kappa_{\max}$ | 2.075 | 0.043 |
| Shape 1 *baseline* | Shape 2 *baseline* | 2.681 | 0.01 |
| Shape 1 *baseline* | Shape 2 $\kappa_{\max, \min}$ | 2.853 | 0.006 |
| Shape 1 *baseline* | Shape 2 $u(\kappa)_{\max}$ | 2.508 | 0.015 |
| Shape 1 *baseline* | Natural shape *baseline* | 2.075 | 0.043 |
| Shape 1 *baseline* | Natural shape $\kappa_{\max}$ | 3.632 | $< .001$ |
| Shape 1 *baseline* | Natural shape $\kappa_{\max, \min}$ | 2.508 | 0.015 |
| Shape 1 *baseline* | Natural shape $\kappa_{\max}$ | 2.681 | 0.01 |
| Shape 1 $u(\kappa)_{\max}$ | Shape 2 $\kappa_{\max, \min}$ | 2.162 | 0.035 |
| Shape 1 $u(\kappa)_{\max}$ | Natural shape $\kappa_{\max}$ | 2.94 | 0.005 |

## Footnotes

1 Curvature can be described in many ways. Here, we define curvature as signed curvature where positive curvature values refer to convex and negative values to concave curvature along the contour (see Method section for formal definition and notation). Hence, curvature *maxima* are convex points on the contour that have the greatest local curvature while curvature *minima* are concave points that have the greatest local curvature.

2 In information theory (Shannon, 1948), the term “information” refers to the reduction of uncertainty about the state of a system or the outcome of a random variable when a particular event is observed. Formally, information is quantified using the concept of surprisal, where rarely occurring events carry more information (i.e., they are *more surprising*) and likely events carry less information (they are *less surprising*). In other words, less probable events are more informative. The formal framework for calculating information along contours that we applied here was developed by Feldman and Singh (2005).

3 Note that the Intersection over Union (IoU, Jaccard index) and the Dice-Sørensen coefficient generate different results, where the Dice-Sørensen coefficient > IoU. However, qualitatively, they produce the same result and interpretation. shapes was equated (see text for rationale). The light-grey shapes in the *Convexities and Concavities* condition illustrate shapes that result from connecting only local concave maxima, which cannot be recognized at all.

4 Any closed shape must contain the same number or more convex points than concavities. Connecting the points of maximum convex curvature represent the outer boundary (hull) of a closed shape, resulting in a polygon that envelops most of the shape. However, polygons created by connecting the concave maxima results in polygons that do not resemble the original closed shape, as convexities tend to extend beyond the polygon’s confines (Schmidtmann et al., 2015). This is illustrated in the form of the grey polygons for the *Convexities and Concavities* condition in Figure 2.

5 We calculated the number of vertices for each condition (1. *Convexities-only,* 2. *Convexities and Concavities,* 3. *Surprisal)* across all animal stimuli. The mean number of points (± standard deviation) were: for 1. 12.2 ±2.3, for 2. 21.6 ±3.9, and for 3. 13.4 ±4.6). Hence, the ratio between the *Surprisal* and *Convexities and Concavities* condition is approximately 0.6 or 60%. In other words, only ∼60% of the high-surprisal points are required to achieve the same performance as in the *Convexities and Concavities* condition.

## Notes

### Competing Interest Statement

The authors have declared no competing interest.

### Summary of Updates

The revised version contains a new Experiment 2 and revised figures.

